# Amelioration of signalling deficits underlying metabolic shortfall in TREM2^R47H^ human iPSC-derived microglia

**DOI:** 10.1101/2024.02.12.579896

**Authors:** Foteini Vasilopoulou, Thomas M. Piers, Wei Jingzhang, John Hardy, Jennifer M. Pocock

## Abstract

Classic immunometabolic switching from oxidative phosphorylation (OXPHOS) to glycolysis occurs in myeloid cells such as microglia when encountering pathogenic insults. We previously showed that patient iPSC-derived microglia (iPS-Mg) harbouring the Alzheimer’s disease (AD) TREM2^R47H^ hypomorph display deficits in the ability to metabolically switch from OXPHOS to glycolysis, with both reduced mitochondrial maximal respiration and glycolytic capacity. To resolve this, we generated common variant, TREM2^R47H^ and TREM2^−/−^ variant human iPS-Mg and we assessed the ability of supplementation with citrate or succinate, key metabolites and cell cycle breaking points upon microglia activation, to overcome deficits in a number of functions we identified previously as affected by TREM2 loss of function phenotypes. Succinate supplementation was more effective than citrate at overcoming mitochondrial deficits in OXPHOS and did not promote a switch to glycolysis. Citrate enhanced the lipid content of TREM2^R47H^ iPS-Mg and was more effective at overcoming phagocytic deficits whereas succinate increased lipid content and phagocytic capacity in TREM2^−/−^ iPS-Mg. Microglia cytokine release upon activation was differentially affected by citrate or succinate and neither metabolite altered soluble TREM2 shedding. Neither citrate nor succinate enhanced glycolysis but drove their effects through oxidative phosphorylation. Our data point to discrete pathway linkage between microglial metabolism and functional outcomes with implications for AD pathogenesis and treatments.

## 1 Introduction

Variants in the triggering receptor expressed on myeloid cells (TREM2) are linked to an increased risk of developing dementia, including late-onset Alzheimer’s disease (AD) (1,2). TREM2 is expressed on microglia, the resident immune cells of the brain, and is required for diverse microglia responses in neurodegeneration, including immunometabolic plasticity, phagocytosis, and survival (3–6). Identifying how TREM2 variants affect the function of microglia is pertinent to our understanding of disease progression in AD and for identifying new pathways for protection.

In a surveillant or homeostatic state, microglia express a reduced protein and pathway activation load. Upon input of appropriate signals, microglia rapidly respond with increased energy production, gene expression, and protein synthesis. In this regard, microglia switch from a dependency on oxidative phosphorylation (OXPHOS) to glycolysis, which, whilst produces less energy (adenosine triphosphate, ATP) compared with OXPHOS, the ATP production rate is faster upon insult (7–9). In TREM2^−/−^ AD animal models, a reduced metabolic ‘fitness’ is associated with TREM2 loss of function(6). Our previous studies in human induced pluripotent stem cell-derived microglia (iPS-Mg) from patients expressing the AD-linked R47H^het^ TREM2 variant and the Nasu Hakola variants T66M^hom^ and W50C^hom^ increased our understanding of the deficits linked to TREM2 loss-of-function (LoF) variants; specifically in aerobic glycolysis and mitochondrial respiration (10–12). Moreover, we reported that iPS-Mg expressing AD-linked TREM2 variants are unable to switch from a mainly OXPHOS state of energy production to one in which glycolysis dominates (11). We also identified that this inability to generate sufficient energy results in changes to the exosome profile of these cells (13,14) and an inability to activate the inflammasome (12).

Recent findings indicate that signalling metabolites play a pivotal role in modulating microglial functions through diverse mechanisms (15–18). At the same time, the activation of immune cells has been associated with tricarboxylic acid (TCA) cycle fragmentation downstream of two key signalling metabolites, namely citrate and succinate (19,20). While citrate can bridge carbohydrate and fatty acid metabolism in the TCA cycle, succinate interacts directly with the mitochondrial electron transport chain (ETC), enabling a ‘shortcut’ route to ATP production via oxidative metabolism. Whilst TREM2 impact on energetic and anabolic pathways has been linked to alterations in glycolytic TCA cycle intermediates, including citrate and succinate, which are reduced in TREM2^−/−^ macrophages (6) the involvement of those energy substrates in TREM2-dependent metabolic reprogramming pathways is still underexplored.

Here we hypothesized that microglia metabolic deficits in TREM2 LoF variants may be linked to metabolite signalling deficits, and in turn, fuelling microglia with these metabolites could restore the functional deficiencies exhibited by TREM2 LoF variant microglia. To address that, we determined the impact of supplementing culture media with citrate or succinate on functions found to be deficient in TREM2 LoF variant human iPS-Mg. Considering that metabolic profile is closely related to the status and functional changes of microglia, this may allow a remedial approach to TREM2 linked deficits. We show that citrate and succinate supplementations affect microglia metabolic processes and restore key functions, providing insights into the metabolic pathways that underly TREM2-related metabolic disturbances in microglia.

## 2 Materials and methods

### 2.1 Human iPSC generation of microglia

Ethical permission for this study was obtained from the National Hospital for Neurology and Neurosurgery and the Institute of Neurology joint research ethics committee (study reference9/H0716/64). The TREM2^R47H^ homozygous line (BIONi010-C7) and TREM2^−/−^ line (BIONi010-C17) were gene edited BioNi010-C, purchased from EBiSC. SFC840 (Stembancc) and BioNi010-C were used as the common variant (Cv) control lines. iPSCs were maintained and routinely passaged in Essential 8 medium. Using our previously described protocol (11), which incorporates procedures from earlier protocols (10,21) iPSC-derived microglia (iPS-Mg) were generated. Experimental replicates were one clone assayed in at least three independent experimental runs (Cv Control, TREM2^R47H^ homozygous and TREM2^−/−^).

### 2.2 Cellular energy analysis

For real-time analysis of oxygen consumption rates (OCR) (using Seahorse XF Cell Mito Stress Test kits), or proton efflux rate (PER) (using Seahorse Glycolytic Rate assays, all Agilent Technologies), iPS-Mg were plated (20 000-25 000/well), matured on Seahorse cell culture microplates and analysed using a Seahorse XFe96 Analyzer (Agilent Technologies) as previously described (11,12). Where indicated, cells were incubated with 10mM succinate (succinic acid, disodium salt, Sigma) or 10mM citrate (sodium citrate, trisodium salt, Sigma) in the presence or absence of 100 ng/ml Lipopolysaccharide (LPS) and 10 U/ml interferon-gamma (IFN-γ) for 24 h prior to analysis. Data were analysed using Wave v2.4.0.6 software (Agilent Technologies).

### 2.3 ELISA analysis of cytokine secretion and shed soluble TREM2

IPS-Mg (50 000/cm^2^) were stimulated with LPS/IFN-γ (100 µg/ml and 10 U/ml respectively) for 24 h in the presence of citrate (10 mM), 2-deoxyglucose (2-DG, 3 mM), citrate plus 2-DG, or succinate (10 mM) alone or with 2-DG for 24 h. Cell culture medium was collected, centrifuged at 300*×g* for 15 min to remove cell debris and typically 50 μl (1:2 dilution) appraised for secreted TNFα or 200 μl (1:2 dilution) for secreted IL-1β using human Quantikine ELISA kits, as per the manufacturer’s instructions (R&D).

Soluble TREM2 (sTREM2) was appraised in 100 μl of cell culture medium with an in house-generated ELISA system as previously described (10,12). Briefly, MaxiSORP 96-well plates were coated with 1μg/ml of a rat anti-mouse/human TREM2 monoclonal antibody (R&D Systems; clone 237920) at 4°C, overnight. After blocking supernatant samples from different groups and standards (recombinant human-TREM2-His; Life Technologies) were incubated for 2 h at room temperature (RT) with biotinylated polyclonal goat anti-human TREM2 capture antibody (0.1 μg/ml; AF1828, R&D Systems). Following incubation with streptavidin-HRP (Invitrogen; 0.1 µg/ml), chromogenic substrate solution was added (TMB, Life technologies) the reaction terminated (with 0.16 M H_2_SO_4_) and absorbance read at 450 nm (EZ Read microplate). Values were normalised to protein content of cell lysates for each sample.

### 2.4 Lipid Spot staining

IPS-Mg (60.000/coverslip) were matured on 13 mm glass coverslips. Cells were treated with citrate (10 mM) or succinate (10 mM) for 24 h and subsequently fixed with 4% PFA in PBS for 20 min at RT. For lipid spot staining, cells were washed with PBS, incubated with Wheat germ agglutinin (WGA) (1:2000) for 5 min, following permeabilisation with 0.1% Triton-X. Cells were then washed with PBS and incubated with LipidSpot^TM^488 (Biotium) (1:1000) for 15 min in the dark. Nuclei were stained with DAPI (1:5000) and coverslips mounted on glass slides using Fluoromount-G (Invitrogen). Three-five different regions per coverslip were imaged on a Zeiss LSM710 confocal microscope using the LSM Pascal 5.0 software.

### 2.5 Lactate release in cell culture medium

Levels of lactate secreted from iPS-Mg into supernatant were determined using an L-lactate Assay kit (abcam) and a D-lactate assay kit (abcam), after 24 h treatment with citrate (10 mM) or succinate (10 mM), and 2-deoxyglucose (2-DG, 3 mM) in the presence or absence of LPS/IFN-γ (100 ng/ml and 10 U/ml respectively) as per the manufacturer’s instructions. Values were normalised to protein content of cell lysates for each sample.

### 2.6 Mitochondrial Superoxide analysis

Mitochondrial superoxide levels were measured by loading iPS-Mg (50.000/cm^2^) with MitoSOX**™** red superoxide indicator (ThermoFisher) and analyzed by flow cytometry as previously described (11). Following 24 h treatments with citrate (10 mM) or succinate (10 mM), cells were incubated with MitoSOX^TM^ red at 5 μM in PBS + 0.5% BSA for 10 minutes at 37°C. 200.000 to 300.000 cells per treatment group were washed and resuspended in PBS followed by flow cytometric analysis (FL2; FACS Calibur, Becton Dickinson). As a positive control for mitochondrial superoxide generation, cells were incubated with rotenone (100 nM) for 30 min before addition of MitoSOX. Data were analyzed using Flowing Software v2.5.2 (University of Turku).

### 2.7 Western blotting

iPS-Mg (50.000/cm^2^) were lysed in RIPA buffer (50 mM Tris, 150 mM NaCl, 1% SDS, and 1% Triton X-100) containing 1x Halt™ protease and phosphatase inhibitor cocktail at 4°C. Lysates were centrifuged at 15,000g for 15 minutes at 4°C to clarify the lysate and separate the soluble and insoluble (nuclear) fractions. Fractions were snap frozen on dry ice and stored at −80°C until use. Soluble fractions were normalised after BCA protein quantification and 4x running buffer (LiCor) plus 10% DTT. Samples were denatured at 95°C for 10 min and run on 4-20% Criterion TGX Precast Midi Protein gels (Bio-Rad). Resolved protein was transferred to nitrocellulose using Trans-Blot turbo transfer packs, washed for 10 minutes at RT with TBS and blocked for 1 hour at RT in 5% milk in TBS-T. Blocked membranes were incubated in primary antibody overnight at 4°C (Phospho-NF-κB p65 (Ser536) (93H1), Cell Signalling, (1:1000); NF-κB p65 (D14E12), Cell Signalling, (1:1000); Anti-β-Actin (AC-15), Sigma Millipore (1:5000); Hsp70 (5A5), abcam, (1:500). Membranes were washed with TBS-T and incubated for 1 hour at RT in the dark with corresponding LiCor compatible Alexafluor 680/800nm conjugated secondary antibodies or HRP-conjugated anti-mouse or anti-rabbit (ECL detection). The membranes were washed 3x with TBS-T followed by a final TBS-T wash and visualised using an Odyssey detection system (LiCor). Protein bands were subsequently quantified using exported zip files in Image Studio Lite v5.2.

### 2.8 Cell stress array analysis

IPS-Mg were incubated for 24 h with citrate (10 mM) or succinate (10 mM) and cell lysates were collected and prepared according to manufacturer’s instructions (Proteome Profiles^TM^ Human cell stress array; Bio-Techne). Total protein quantification was performed on the aliquots of each treatment group for data normalization purposes. Cell lysates from 3-4 independent experiments were pooled according to the TREM2 genotype after basal or citrate/succinate treatment. Data were analyzed and quantified using exported zip files in Image Studio Lite v5.2. Data were plotted as relative protein expression, normalized to total cellular protein levels.

### 2.9 Aβ_1-42_ (HiLyte488) phagocytosis

IPS-Mg were plated at a density of 50.000/cm^2^ and were pre-treated with citrate (10 mM), succinate (10 mM), and 2-DG (3 mM) for 24 h. Oligomerised Aβ_1-42_ (HiLyte488; 100 nM) was added to all groups and cells were incubated at 37°C + 5% CO_2_ for 2 h. Cytochalasin-D (CytoD; 100 µM) was added to negative control groups for 30 min prior to Aβ_1-42_ addition. 200.000 to 300.000 cells were then collected, centrifuged at 300g for 3min at RT and the cell pellets were resuspended in PBS for FACS analysis (FL1; FACs Calibur, Beckton Dickinson). Data were analysed using Flowing software v2.5.1 (Turku Centre of Biotechnology, University of Turku).

### 2.10 Data analyses and statistics

All experiments were performed on at least 3 independent cell plantings with technical replicates of at least 3 per experiment. Where indicated data were normalized to cell protein content (BCA) or cell number (Crystal violet). For cell proteome arrays were analysed on pooled samples from three different cell platings. Data are presented as mean ± SEM (n = 3 - 12). Statistical significance was addressed using one-way ANOVA or two-way ANOVA with Tukey post-hoc analysis to compare controls and treated groups respectively unless otherwise stated. Statistical significance was considered when p-values were <0.05. Statistical outliers were determined with Grubbs’ test and, when necessary, were removed.

## 3 Results

### 3.1 TREM2 variant deficits in maximal respiration and spare respiratory capacity are rescued with succinate but not citrate (Figure 1)

We performed seahorse metabolic analysis of oxidative phosphorylation (OCR) in iPS-Mg expressing TREM2 Cv, TREM2^R47H^ or TREM2^−/−^ after a 24 h preincubation with media supplemented with succinate (10 mM) (**Figure 1A**) or citrate (10 mM) (**Figure 1B**). The results indicated that deficits in maximal respiration and spare respiratory capacity exhibited in iPS-Mg expressing the TREM2^R47H^ were rescued when cells were pre-incubated with succinate but not with citrate (**Figure 1C-E**). In TREM2^−/−^ iPS-Mg, only succinate supplementation further increased these parameters (**Figure 1C-E**). None of the substrates significantly affected the respiration in Cv iPS-Mg (**Figure 1C and 2B**). LPS/IFN-γ stimulation for 24 h increased maximal respiration in all iPS-Mg lines tested (Supplementary figure S1).

**Figure 1:**
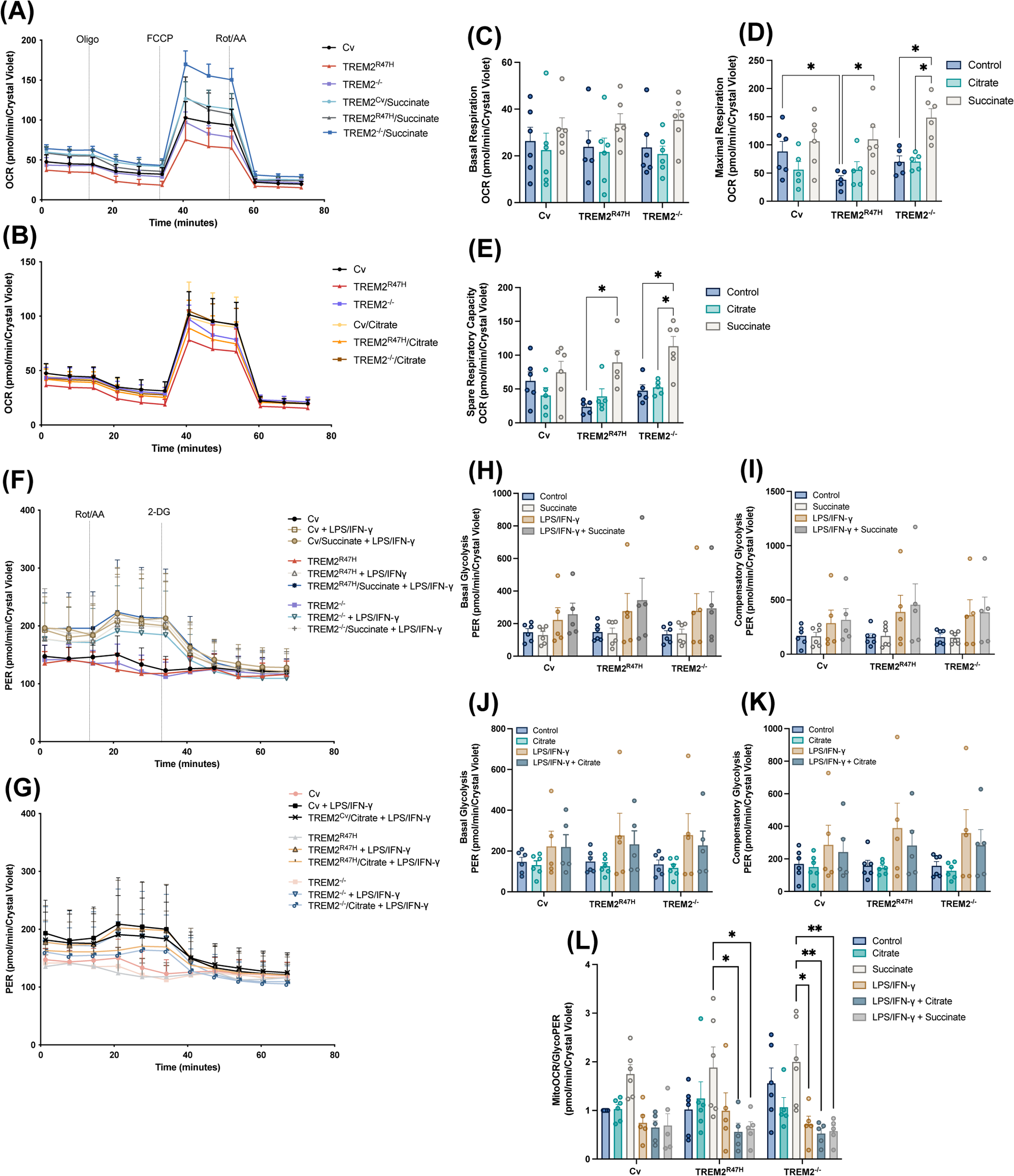
Metabolic function of iPS-Mg supplemented with citrate or succinate. **(A, B)** Metabolic analysis of oxygen consumption rate (OCR) and **(F, G)** proton efflux rate (PER) in Common variant (Cv), TREM2^R47H^ and TREM2^−/−^ iPS-Mg after 24 h treatment with succinate (10 mM) or citrate (10 mM)**. (C)** Basal respiration, **(D)** maximal respiration, **(E)** spare respiratory capacity**, (H, J)** basal glycolysis, **(I, K)** compensatory glycolysis, and **(L)** mitoOCR/GlycoPER ratio in basal conditions and upon LPS/IFNγ stimulation in iPS-Mg expressing Cv, TREM2^R47H^ or TREM2^−/−^ after incubations with citrate or succinate. Data are presented as mean ± SEM. Statistical significance was addressed using One-way ANOVA or Two-way ANOVA with Tukey post-hoc analysis to compare controls and treated groups respectively; *p<0.05 (n=5-6)

**Figure 2:**
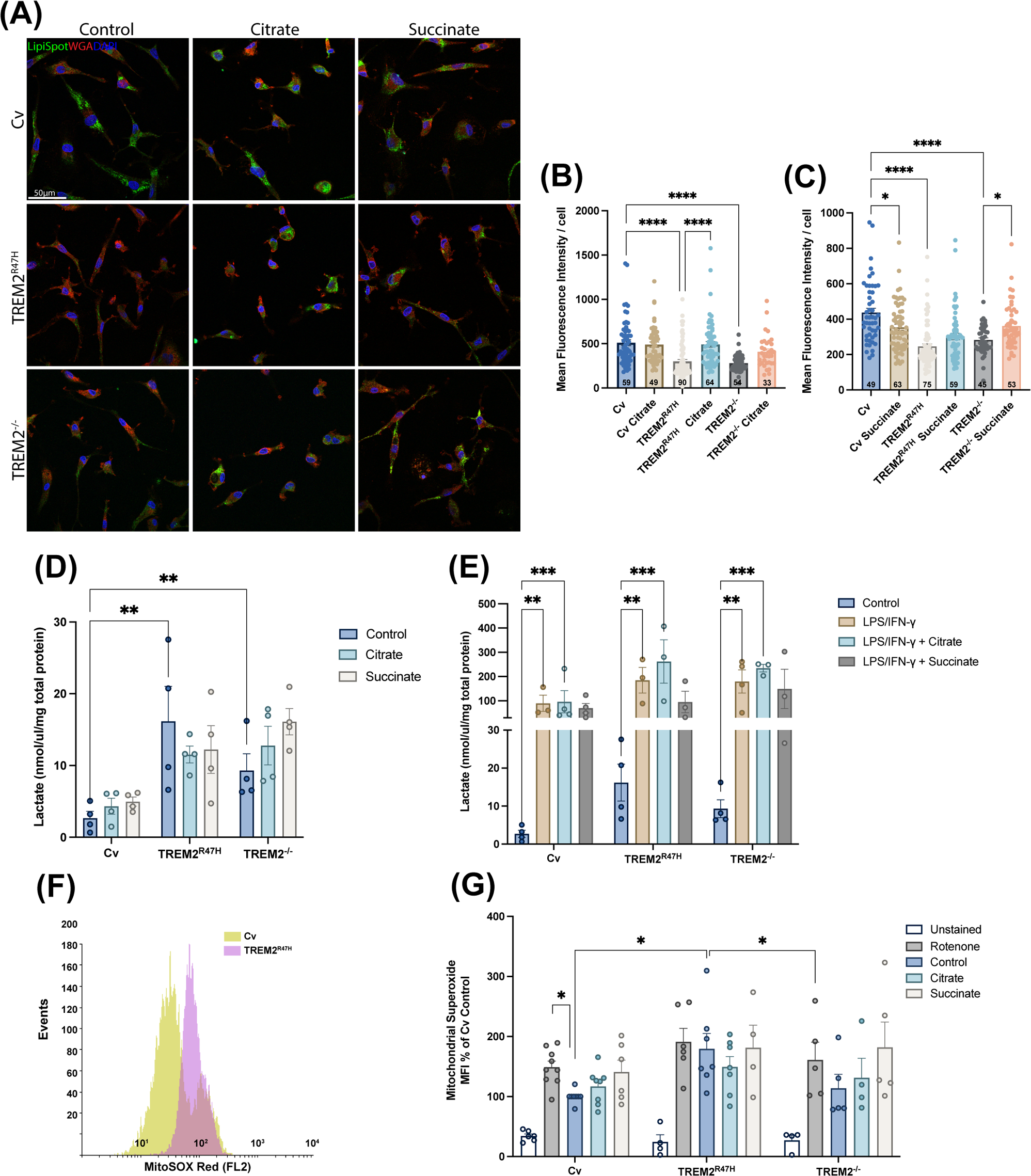
Lipid content, lactate and mitochondrial superoxide production in iPS-Mg supplemented with citrate or succinate. **(A)** Representative images and (**B, C**) quantification of LipidSpot^TM^488 staining of lipid droplets in Common variant (Cv), TREM2^R47H^ and TREM2^−/−^ variant iPS-Mg after medium supplementation with citrate (10mM) or succinate (10mM) for 24h. **(D)** Basal lactate production and **(E)** lactate production upon LPS/IFN-γ stimulation were determined in the iPS-Mg cell culture medium after supplementation with citrate or succinate. **(F)** Representative histogram and **(G)** mitochondrial superoxide (MitoSOX) production as determined by FACS in Cv, TREM2^R47H^ and TREM2^−/−^ variant iPS-Mg after succinate or citrate supplementation or rotenone treatment (positive control). Data are presented as mean ± SEM. Statistical significance was addressed using One-way ANOVA or Two-way ANOVA with Tukey post-hoc analysis to compare controls and treated groups respectively; *p<0.05 (n=5-6)

### 3.2 Compensatory glycolysis is not affected by citrate or succinate (Figure 1)

Metabolic analyses of the glycolytic rate (PER) in iPS-Mg with Cv, TREM2^R47H^ or TREM2^−/−^ (**Figure 1F-G**) showed that after 24 h preincubation, LPS/IFN-γ promoted a swift increase in glycolysis as shown by a decrease in MitoOCR/GlycoPER ratio in all iPS-Mg genotypes (**Figure 1L**). Succinate or citrate treatments, however, did not affect basal glycolysis (**Figure 1H and 1J**) or compensatory glycolysis (**Figure 1I and 1K**).

### 3.3 Lipid deficits are differentially modified upon treatment with citrate and succinate (Figure 2)

TREM2 is a modulator of lipid metabolism (22). To determine the influence of citrate or succinate on the lipid content of iPS-Mg with TREM2 variants, we analysed lipid droplet content in these cells by Lipid Spot staining (**Figure 2A**). TREM2^R47H^ or TREM2^−/−^ iPS-Mg showed a significant reduction in overall lipid droplet content/cell compared with Cv iPS-Mg (**Figure 2B and 2C**). Treatment with citrate (10 mM) did not significantly affect the lipid content/cell of Cv or TREM2^−/−^ iPS-Mg (**Figure 2B**) but significantly enhanced the lipid content in TREM2^R47H^ iPS-Mg. Succinate produced a significant but small drop in lipid drop content in Cv, did not significantly affect it in TREM2^R47H^, but produced a slight but significant increase in TREM2^−/−^ iPS-Mg (**Figure 2C**).

### 3.4 Lactate metabolism is enhanced with LPS/IFN-γ treatment but not modified with citrate or succinate (Figure 2)

As lactate is a waste product of cells with high metabolic rate and is implicated in immunometabolism of microglia (23) as well as neurodegeneration (24), the secretion of both lactate enantiomers from iPS-Mg was investigated, since both D-lactate and L-lactate are produced by microglia (24). Interestingly the levels of L-lactate secreted were different across the TREM2 variants at basal levels, with more lactate secreted by TREM^R47H^ and TREM2^−/−^ iPS-Mg compared with Cv iPS-Mg (**Figure 2D**), and significantly enhanced in all variants when the cells were incubated with LPS/IFN-γ for 24 h (**Figure 2E**). However, neither citrate nor succinate significantly altered L-lactate secretion across any of the iPS-Mg lines (**Figure 2D and 2E**). D-lactate was not secreted at basal conditions to detectable levels (data not shown).

### 3.5 Mitochondrial superoxide production is not modulated by citrate or succinate (Figure 2)

We showed previously that iPS-Mg expressing TREM2^R47H^ heterozygous produced significantly more mitochondrial-generated superoxide than TREM2 Cv expressing microglia at basal levels (11). This was also the case here for TREM2^R47H^ homozygous compared with Cv (**Figure 2F and 2G**) whereas superoxide production increase in TREM2^−/−^ iPS-Mg was slight but not significant (**Figure 2G**). Media supplementation with citrate or succinate for 24 h did not significantly modify superoxide production across any of the lines (**Figure 2G**).

### 3.6 Deficits in Aβ_1-42_ uptake are restored by citrate (Figure 3)

Previously we and others have identified an aberrant phagocytic phenotype in iPS-Mg expressing TREM2 LoF variants, both in their ability to phagocytose apoptotic cells (25) and in their ability to phagocytose Aβ_1-42_ (11). Here preincubation with citrate but not succinate was able to restore the impaired Aβ_1-42_ phagocytosis in TREM2^R47H^ iPS-Mg (**Figure 3A and 3B**), and both metabolites increased Aβ_1-42_ uptake in TREM2^−/−^ iPS-Mg (**Figure 3B**). Inhibition of glycolysis by 2-DG led to impaired Aβ_1-42_ phagocytosis in Cv iPS-Mg which was restored by citrate but not by succinate (**Figure 2C**). Furthermore, citrate was unable to reverse the effects of glycolytic inhibition by 2-DG in TREM2^R47H^ and TREM2^−/−^ iPS-Mg and succinate had no effect (**Figure 2C**).

**Figure 3:**
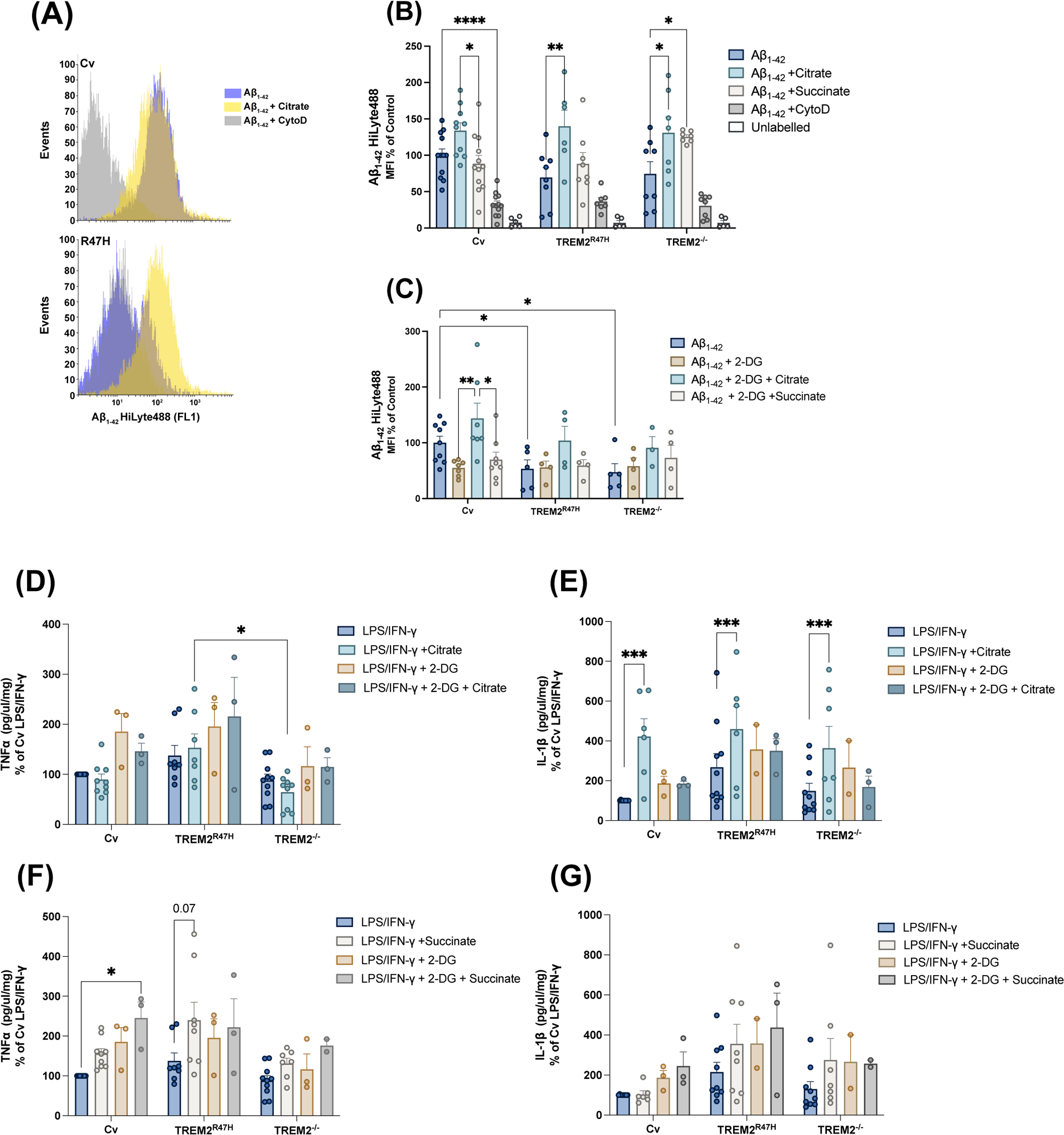
Effect of citrate and succinate on Aβ_1-42_ phagocytosis and cytokine secretion by iPS-Mg. **(A)** Representative histograms of Aβ_1-42_ uptake in iPS-Mg upon citrate supplementation. **(B)** The ability of common variant (Cv), TREM2^R47H^ and/or TREM2^−/−^ variant iPS-Mg to phagocytose Aβ_1-42_ oligomers was assessed after citrate and succinate supplementation in basal conditions and **(C)** upon inhibition of glycolysis by 2-Deoxyglucose (2-DG). **(D, E)** LPS/IFN-γ evoked pro-inflammatory release of TNFα and **(F, G)** IL-1β were assessed by ELISA in the culture medium of iPS-Mg treated with citrate or succinate in basal conditions and after glycolytic inhibition by 2-DG. Data are presented as mean ± SEM. Statistical significance was addressed using One-way ANOVA or Two-way ANOVA with Tukey post-hoc analysis to compare controls and treated groups respectively; *p<0.05 (n=2-12)

### 3.7 Inflammatory cytokine secretion is differentially modified by citrate and succinate (Figure 3)

Since the secretion of inflammatory cytokines is implicated in AD (26,27) and at the same time metabolic intermediates can act as pro-inflammatory agents per se (15,28–30), we investigated whether TNFα or IL-1β secretion is modulated by medium supplementation with succinate or citrate (**Figure 3D-G**). Basal levels of TNFα or IL-1β were very low (**Supplementary figure S2**) and were not altered by citrate or succinate (data not shown), thus we investigated the modulation of evoked cytokine secretion. Citrate did not significantly modify TNFα secretion evoked with LPS/IFN-γ in any of the lines or following co-treatment with 2-DG compared to untreated cells (**Figure 3D**). However, TNFα secretion was significantly lower in TREM2^−/−^ compared to TREM2^R47H^ upon citrate supplementation (**Figure 3D**). Furthermore, citrate significantly enhanced LPS/IFN-γ evoked IL-1β secretion, but not that of 2-DG (**Figure 3E**). Succinate increased LPS/IFN-γ evoked TNFα secretion in Cv iPS-Mg which was even higher upon 2-DG inhibition (**Figure 3F**). Similarly, succinate elevated secretion in TREM2^R47H^ iPS-Mg without blocking glycolysis with 2-DG (**Figure 3F**). TREM2^−/−^ lines showed no significant change in TNFα secretion with succinate supplementation in any condition (**Figure 3G**). There was no significant effect of succinate on IL-1β secretion in any treatment or TREM2 line compared with LPS/IFN-γ alone (**Figure 3G**).

### 3.8 Soluble TREM2 shedding is not reduced by citrate or succinate (Figure 4)

Since sTREM2 shedding is reduced in TREM2^R47H^ variants, we investigated the effects of citrate or succinate on this iPS-Mg variant as well as on Cv. TREM2^−/−^ iPS-Mg served as negative controls for sTREM2 shedding. As expected, sTREM2 secretion was significantly reduced in Cv and TREM^R47H^ following LPS/IFN-γ stimulation compared with basal levels (**Figure 4A**). Furthermore, less sTREM2 was shed at basal or following stimulation in TREM2^R47H^ compared with TREM2^Cv^ iPS-Mg (**Figure 4A**). Citrate did not significantly modify the release of sTREM2 in either Cv or R47H expressing iPS-Mg at basal or evoked conditions (**Figure 4A**). Succinate did not modify basal sTREM2 secretion or the reduction evoked with LPS/IFN-γ stimulation across any of the lines (**Figure 4A**).

**Figure 4:**
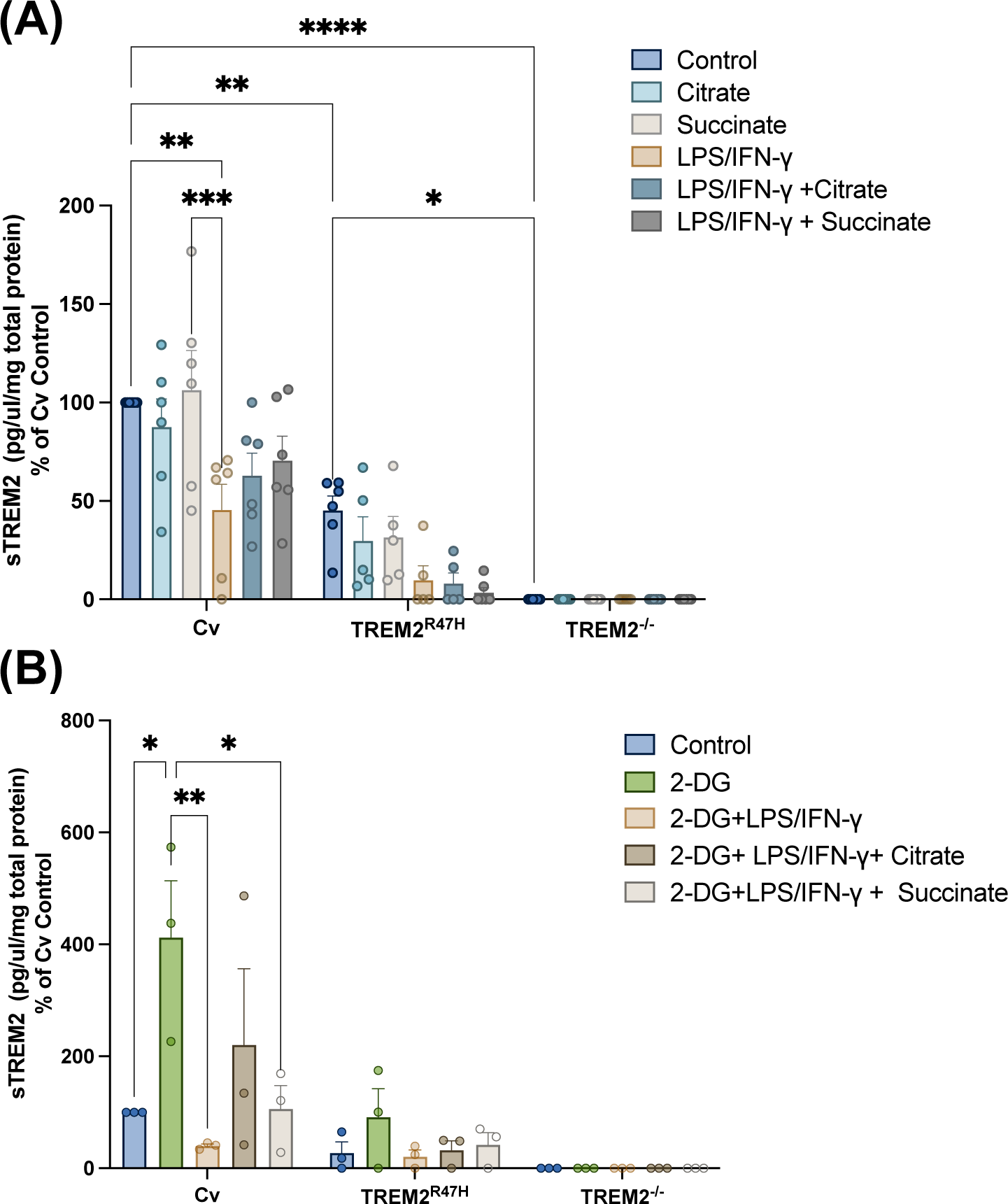
Effect of citrate or succinate on shed TREM2 in iPS-Mg in basal conditions and upon microglia stimulation. **(A)** Shed soluble TREM2 (sTREM2) was determined in the culture medium of unstimulated common variant (Cv), TREM2^R47H^ and TREM2^−/−^ iPS-Mg after treatments with citrate or succinate and/or **(B)** 2-DG mediated inhibition of glycolysis in basal conditions and upon LPS/IFNγ stimulation. Data are presented as mean ± SEM. Statistical significance was addressed using Two-way ANOVA with Tukey post-hoc analysis; *p<0.05 (n=3-8)

Inhibition of glycolysis by 2-DG led to increased basal sTREM2 shedding in iPS-Mg but not in TREM2^R47H^ (**Figure 4B**). LPS/IFN-γ evoked shedding in the TREM2 Cv iPS-Mg was significantly reduced by 2-DG and interestingly this was partially reversed by the presence of citrate (**Figure 4B**). Neither citrate nor 2-DG significantly modified basal or evoked sTREM2 shedding in TREM2^R47H^ iPS-Mg. Similar findings were observed with succinate (**Figure 4B)**.

### 3.9 Stress responses in TREM2 variants are reduced at basal in Cv by citrate or succinate but not modified in R47H or TREM2^−/−^ (Figure 5)

To investigate the effects of citrate or succinate on cell stress pathways TREM2 LoF iPS-Mg we used cellular stress proteome arrays (**Figure 5A-D**). The total protein expression changes of stress proteins in Cv iPS-Mg was reduced by both citrate and succinate followed the same trend in TREM2 ^R47H^ but this was ineffective in TREM2^−/−^ lines (Figure 5A-5D). To investigate this further we selected two proteins which showed high expression in all the iPS-Mg, namely 70 kDa heat-shock protein (HSP70) and Nuclear factor kappa B (NFκΒ), both of which are implicated in AD and metabolic pathways (31,32), and analysed changes in their expression in the presence of citrate or succinate at basal or following LPS/IFN-γ stimulation. Levels of expression of HSP70 were not significantly elevated in Cv iPS-Mg in response to citrate, succinate alone (**Figure 5E and 5I**) or in concert with LPS/IFN-γ stimulation (**Figure 5G and 5I**). However, the level of HSP70 was significantly elevated at basal in TREM2^R47H^ iPS-Mg compared with TREM2^Cv^ iPS-Mg but not modified by citrate or succinate or enhanced significantly by LPS/IFN-γ (**Figure 5G and 5I**). Interestingly there was no elevation at basal in TREM2^−/−^ iPS-Mg compared with Cv iPS-Mg (**Figure 5G and 5I**). The effect of citrate or succinate on NFκB activation was measured as the ratio of p-NFkB to the total NFkB levels on iPS microglia lysates in the absence (**Figure 5E and 5I**) or presence (**Figure 5H and 5I**) of LPS/IFN-γ treatment. No significant differences were observed upon treatments or among the lines.

**Figure 5:**
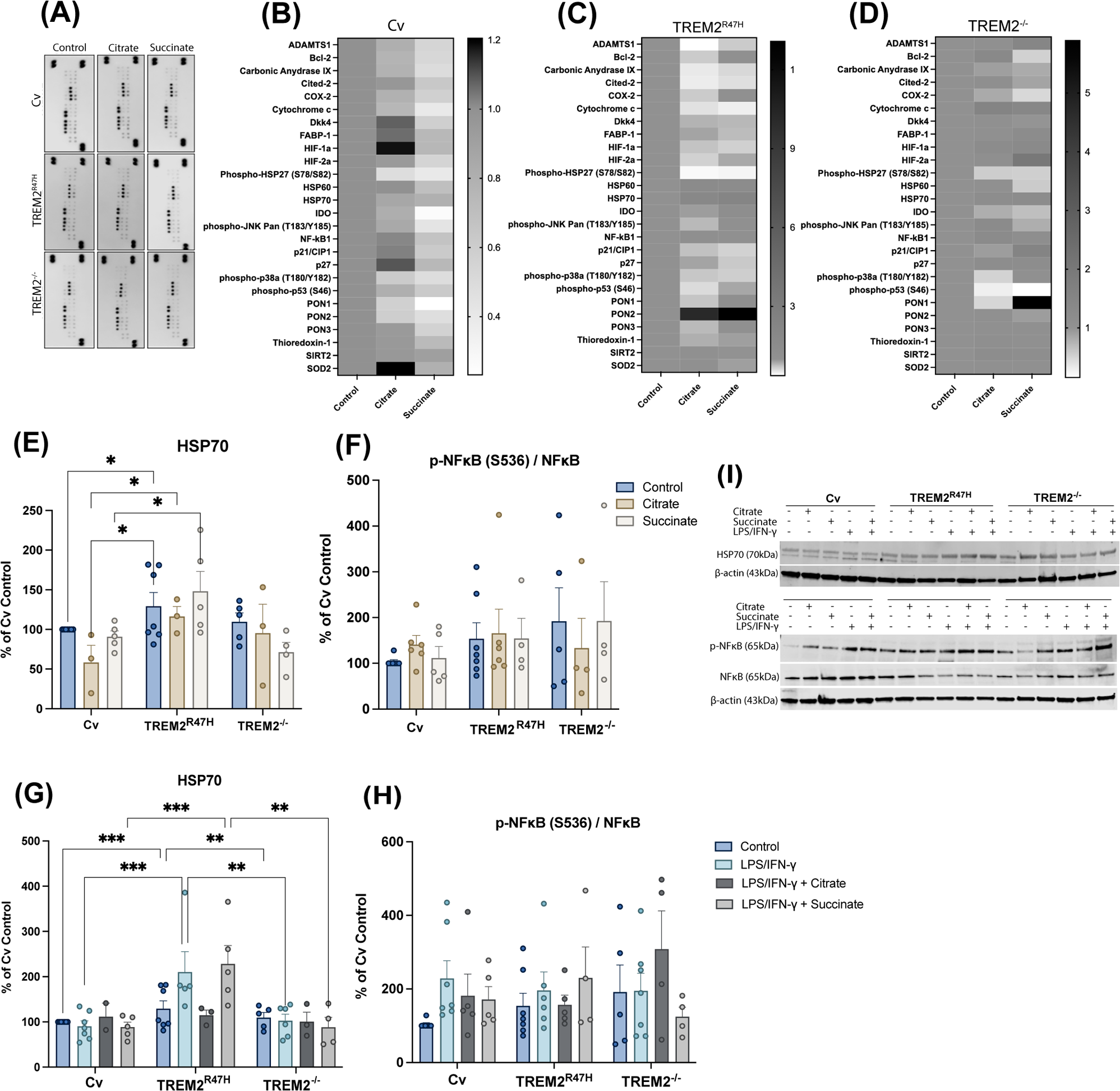
Cellular stress proteome of iPS-Mg is influenced by citrate or succinate. **(A)** Representative cellular stress proteome array dot blots and heatmaps from **(B)** Common variant (Cv), **(C)** TREM2^R47H^, **(D)** TREM2^−/−^ variant iPS-Mg lysates after 24h incubations with citrate (10mM) or succinate (10mM). **(I)** Representative western blots and quantifications for HSP70 **(E)** at basal and **(G)** upon LPS/IFN-γ stimulation, and ratio p-NFκB/NFκB **(F)** at basal and **(H)** following LPS/IFN-γ treatment, in Cv, TREM2^R47H^ and TREM2^−/−^ variant iPS-Mg after treatments with citrate or succinate. Citrate supplemented Cv iPS-Mg group is not included in the selected representative blots **(I**). Data are presented as mean ± SEM. Statistical significance was addressed using Two-way ANOVA with Tukey post-hoc analysis; *p<0.05.

## 4 Discussion

Metabolic sensing in microglia and other immune cells is an important control of their ability to respond to their surroundings. Deficits in lipid sensing and metabolism are associated with TREM2 LoF variants in microglia (10,11,22,33,34), and TREM2 deficiency contributes to exacerbating AD pathology *in vivo* by impairing autophagic and metabolic processes (35). At the same time, TREM2 deficiency has been associated with decreased glycolytic and TCA metabolites *in vitro* (11,35) suggesting that metabolic and functional deficits displayed by TREM2 LoF variant microglia are attributable to deficits in TCA-intermediate signalling.

Citrate and succinate are key signalling TCA intermediates and cell cycle breaking points upon insult (18,36,37). Their levels dynamically change during microglia metabolic reprogramming in a disease context-dependent manner, affecting microglia metabolic status and functions (19,38,39). Here, we investigated whether fueling TREM2 LoF variant iPS-Mg with citrate or succinate could restore metabolic deficits. We show that citrate is not as effective as succinate at enhancing impaired maximal respiration and spare respiratory capacity in TREM2^R47H^ and TREM2^−/−^ iPS-Mg, whilst neither of these substrates significantly modified respiration of Cv iPS-Mg or affected or promoted a glycolytic switch in any of the lines. Previous studies have demonstrated the ability of succinate to boost oxidative phosphorylation in diverse models of diseases characterized by energy dysfunction (40,41). Indeed, succinate is an important OXPHOS component that acts synergistically with NADP for electron supply of different complexes of the respiratory chains (42); thus, less efficient OXPHOS in TREM2^R47H^ and TREM2^−/−^ iPS-Mg may be linked to either impaired succinate signalling or substrate deficiency in microglia, as it has been reported for macrophages (35) Whilst both citrate and succinate have been involved in ROS generation or elimination and glial mitochondria function (28,29,38,43) none of the metabolites appeared to alter mitoSOX levels which were higher in TREM2^R47H^ iPS-Mg reflecting mitochondria abnormalities consistent with our previous findings (11). Notably, disodium succinate, which we also used in this study as an activator of the extracellular pathway, was reported to mediate no effect on ROS levels or mitochondrial fission in primary microglia in contrast to the cell-permeable diethyl succinate (15). Thus, we cannot rule out potential succinate receptor (SUCNR1)-independent effects on TREM2 LoF related mitochondria abnormalities. Alterations observed here in proteins involved in mitochondrial stress pathways such as superoxide dismutase 2 (SOD2), or hypoxia inducible factor 1 α (HIF1α) upon succinate or citrate supplementation that we detected across the lines in the proteome array, are suggestive of an impact on mitochondria function possibly attributable to their impact on OXPHOS that is perturbed in TREM2 LoF variants. Meanwhile, studies have pointed out that it is the shift in microglial metabolism that draws a link between TREM2 deficiency and a disease-associated microglia (DAM) profile. We show that citrate and succinate supplementation of microglia led to an overall decrease of the cell stress proteome, suggesting that TCA metabolites may be beneficial for microglia and compensate for energy deficits due to TREM2 loss of function. Indeed, succinate supplementation was shown to be protective in a mixed microglia culture model of mitochondria dysfunction and in TBI patients (40,41).

TREM2 is a modulator of lipid metabolism, and the R47H variant of TREM2 has reduced affinity for apolipoproteins (e.g., apolipoprotein E; APOE) or lipid ligands, which may contribute to the increased late onset AD risk associated with this TREM2 variant (22,34). This has been suggested as a consequence of the inability of the microglia to process lipid sufficiently, leading to chronic lipid accumulation, particularly of myelin and APOE (5,44–48). Our study shows that TREM2^R47H^ and TREM2^−/−^ iPS-Mg displayed an overall lower lipid accumulation under basal conditions, but when treated with citrate, and to a lesser extent with succinate, the lipid levels increased to those of Cv. These findings suggest that driving OXPHOS in these cells promotes lipid accumulation to levels observed in Cv cells. Furthermore, it is substantiated that increased citrate availability leads to FA synthesis and lipogenesis, which, although detrimental in healthy conditions, is disturbed in neurodegenerative diseases (29). This is further corroborated by the observed effect of citrate and succinate on metabolically healthy iPS-Mg, mediating either no change (citrate) or decreased (succinate) lipids levels in contrast with TREM2 LoF iPS-Mg. Although it has been proposed that chronic lipid accumulation occurs in models of TREM2^−/−^ (34,49), recent evidence suggests a downregulation of lipid metabolism in microglia expressing the TREM2 LoF pQ33X mutation associated with Nasu-Hakola disease (50) in accord with our results. We previously found that exposure of TREM2^R47H^ variant microglia to cells expressing phosphatidylserine (PS+), ameliorated some of the mitochondrial deficits we reported (11,12), and we suggested that where lipid signalling is used to compensate for lack of energy, the R47H variant may prevent ‘efficient’ use of a lipid energy source.

In line with the lack of apparent effects on glycolysis delivered by citrate or succinate and due to their ability to primarily influence OXPHOS, supplementation did not affect the production of the glycolytic by-product, lactate, in our model. However, extracellular lactate levels were markedly increased upon LPS/IFN-γ induced microglia activation, consistent with previous *in vitro* findings (51,52). During glycolysis, pyruvate is turned into the TCA by-product lactate anaerobically, stimulated by PI3K/Akt signalling. However, no differences in the detected lactate levels upon supplementation cannot be interpreted as lack of effect on this glycolytic process. In fact, previous studies show that succinate supplementation altered the lactate/pyruvate ratio, a clinical marker of tissue cerebral metabolic state in disease, *in vivo* and *in vitro,* without significant differences in lactate concentration following succinate treatments (40,41). It is also important to note that lactate release was higher in TREM2^R47H^ and TREM2^−/−^ lines when compared with Cv iPS-Mg in basal conditions and upon LPS/IFN-γ stimulation. It is, therefore, likely that the reported glycolytic deficits of TREM2 deficient microglia are associated with disruptions during glucose conversion into ATP through pyruvate production that result in lactate accumulation. Recently, increased lactate levels were reported in the microglia of AD mouse models characterized by amyloidosis, and a glycolysis/H412 lactylation/PKM2 positive feedback loop in microglia that exacerbates glucose metabolism disorder and proinflammatory activation of microglia in AD (53). These results point to a role of TREM2 signalling in microglial lactate metabolism which deserves further exploration.

To understand further the functional consequences of energy substrate supplementation, we analysed the ability of the cells to phagocytose pathogenic insults such as Aβ_1-42_ and secrete cytokines, functions which we have previously shown to be impaired in R47H variant iPS-Mg (11). Here, citrate and succinate supplementation restored the deficient phagocytic capacity in TREM2^R47H^ (citrate) or TREM2^−/−^ (citrate, succinate), suggesting that boosting OXPHOS (succinate) and/or lipid metabolism (citrate, succinate) was able to compensate for energy deficits in the cells. Upon inhibition of glycolysis, we showed reduced Aβ_1-42_ phagocytosis by Cv iPS-Mg. Conversely, the reduced phagocytic capacity of TREM2^R47H^ and TREM2^−/−^ iPS-Mg was unaffected by 2-DG inhibition, demonstrating Cv iPS-Mg primary dependency on glycolysis to respond to pathology which is not the case for the TREM2 LoF variant microglia. In this setting, citrate reversed 2-DG induced phagocytic impairments in Cv iPS-Mg, further supporting the hypothesis that substrate supplementation can supply the cells with the necessary energy upon metabolic stress. The effect of the substrates on the pro-inflammatory response was less evident even though citrate and succinate have been reported to be pro-inflammatory agents per se, along with existing evidence suggesting a pro-inflammatory response upon activation of the succinate receptor (SUCNR1) in the presence of extracellular succinate (15,28,29). Whilst citrate or succinate supplementation did not affect iPS-Mg basal cytokine secretion to detectable levels, both substrates promoted a subtle pro-inflammatory microglia response following LPS/IFN-γ stimulation. Succinate has been shown to lead to sustained IL-1β production through HIF1-α stabilization (28). Our data point to larger effects of succinate and citrate on cytokine secretion by TREM2^R47H^ and TREM2^−/−^ cells than Cv, and are in line with the observed effects of the substrates on OXPHOS. Upon inhibition of glycolysis by 2-DG and LPS/IFN-γ-evoked TNFα, and IL-1β secretion did not decrease as previous studies have reported (54,55), but instead, their secretion was further amplified in the presence of substrates, suggesting that alternative pathways other than the glycolysis may also drive pro-inflammatory cytokine secretion in the cells. There is significant interplay between cytokines and glucose/lipid metabolism (56), regarding the influence of cytokines on these processes. Altered glucose metabolism can affect cytokine production from human peripheral blood mononuclear cells (PBMCs) and the human monocyte cell line, THP-1 (57).

TREM2 undergoes ectodomain shedding by ADAM10/17 proteases, resulting in of soluble sTREM2 shedding. In AD, sTREM2 has been suggested to either indicate reduced TREM2 signalling and thus impaired microglia functions, or to be protective (58). The relationship between sTREM2 release and metabolic fitness is poorly explored. First, we found that LPS/IFN-γ resulted in decreased sTREM2 shedding in culture supernatant, suggesting that TREM2 shedding is decreased upon LPS/IFN-γ-induced glycolytic switch. However, given that pro-inflammatory conditions such as LPS/IFN-γ stimulation have been previously shown to reduce TREM2 expression resulting in decreased sTREM2 release and detection (59), we additionally evaluated sTREM2 shedding after blocking glycolysis with 2-DG and found its levels accordingly increased. These results show for the first time that immunometabolic processes may influence TREM2 processing, whilst this did not seem to be affected by citrate or succinate supplementation in basal conditions. Upon concomitant 2-DG and LPS/IFN-γ stimulation, sTREM2 levels tended to be increased by citrate and to a lesser extent by succinate suggesting that alternative sources of energy provided by substrate fueling of the cells upon stress insults influence the sTREM2 shedding process in Cv iPS-Mg. Similar trends were observed in different conditions in TREM^R47H^ iPS-Mg, which further support this hypothesis even though they did not reach significance, likely due to the already low basal sTREM2 levels in TREM2^R47H^ lines. Considering that recent data highlight the role of sTREM2 not only in microglial functions but also in microglia-neuronal interactions in AD (60,61) understanding the link between sTREM2 and immunometabolism could provide new insights into the pathogenesis of AD to support the development of new therapies.

Our results show that TCA metabolite signalling pathways are perturbed in TREM2 LoF variant iPS-Mg contributing to the observed metabolic deficits displayed by those cells **(Figure 6**), providing insights into the relationship between dysfunctional metabolism and TREM2 with implications for AD pathogenesis and treatments.

**Figure 6:**
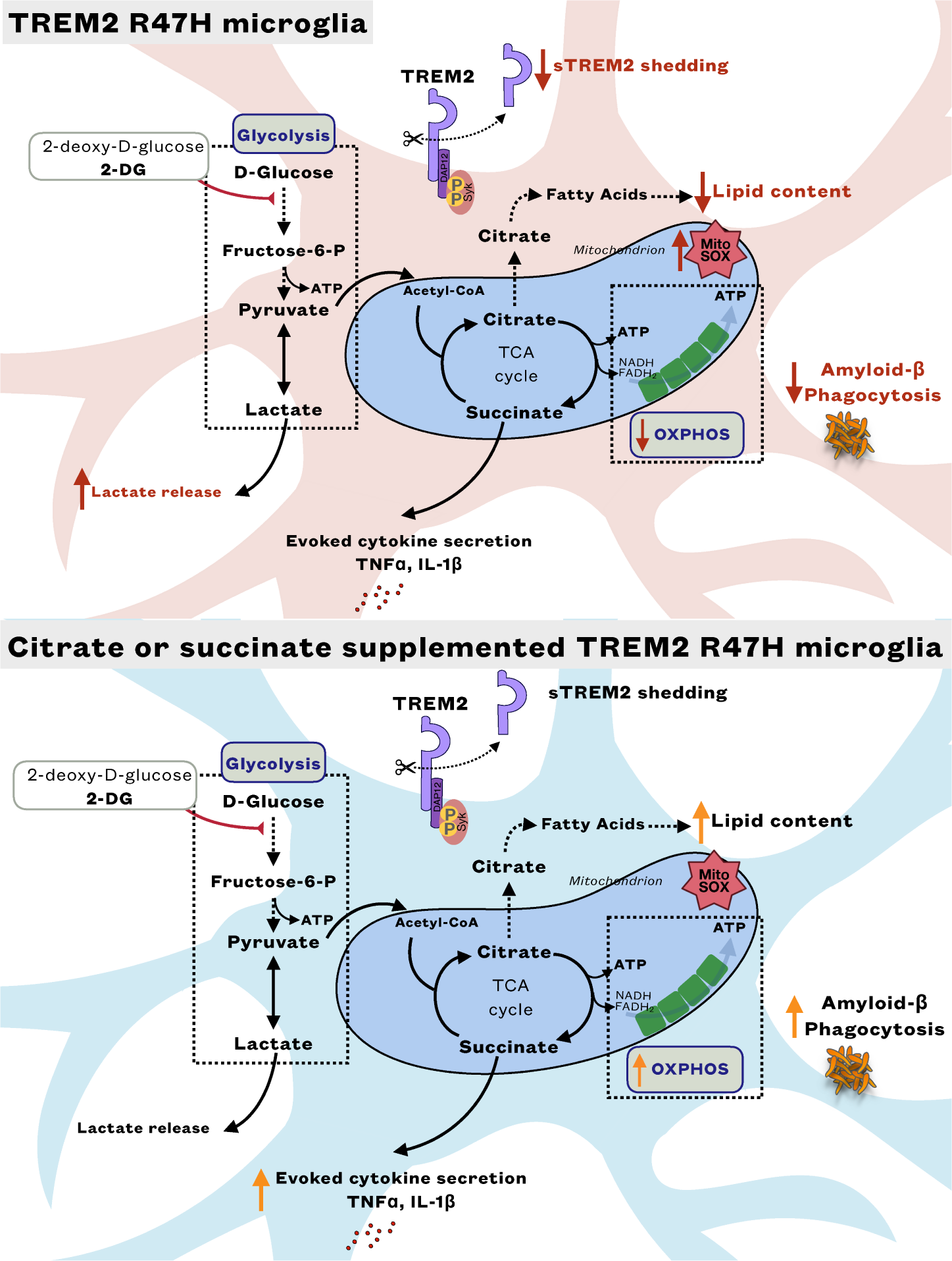
Graphical summary of changes in TREM2^R47H^ iPS-Mg after supplementation with citrate or succinate.

## Abbreviations

2-DG: 2-deoxyglucose
ATP: adenosine triphosphate
AD: Alzheimer’s disease
APOE: apolipoprotein E
CV: Common variant
DAM: disease-associated microglia
ECT: electron transport chain
HIF1α: hypoxia inducible factor 1 α
iPS-Mg: induced pluripotent stem cell-derived microglia
LoF: loss-of-function
OXPHOS: oxidative phosphorylation
OCR: oxygen consumption rates
PER: proton efflux rate
sTREM2: soluble TREM2
SOD2: superoxide dismutase 2
TCA: tricarboxylic acid
triggering receptor expressed on myeloid cells: TREM2.

## 5 Conflict of Interest

The authors declare that the research was conducted in the absence of any commercial or financial relationships that could be construed as a potential conflict of interest.

## 6 Authors Contributions

J.M.P, T.M.P and F.V. designed the study, F.V., and T.M.P., carried out the experiments and analysed the data, J.Wei performed additional data analyses, J.M.P., F.V. and T.M.P. wrote the paper. All authors provided comments, contributed to and agreed on the final version of the paper.

## 7 Funding

F. Vasilopoulou and T.M. Piers and were supported by funding to J.M. Pocock and J. Hardy from the Innovative Medicines Initiative 2 Joint Undertaking under grant agreement No.115976, “PHAGO”. This Joint Undertaking receives support from the European Union’s Horizon 2020 research and innovation programme and EFPIA. This work was supported by the Medical Research Council Core funding to the MRC LMCB (MC_U12266B) and Dementia Platform UK (MR/M02492X/1).

## 8 Data Availability

All data are available on request.

---

**Supplementary Figure 1.**
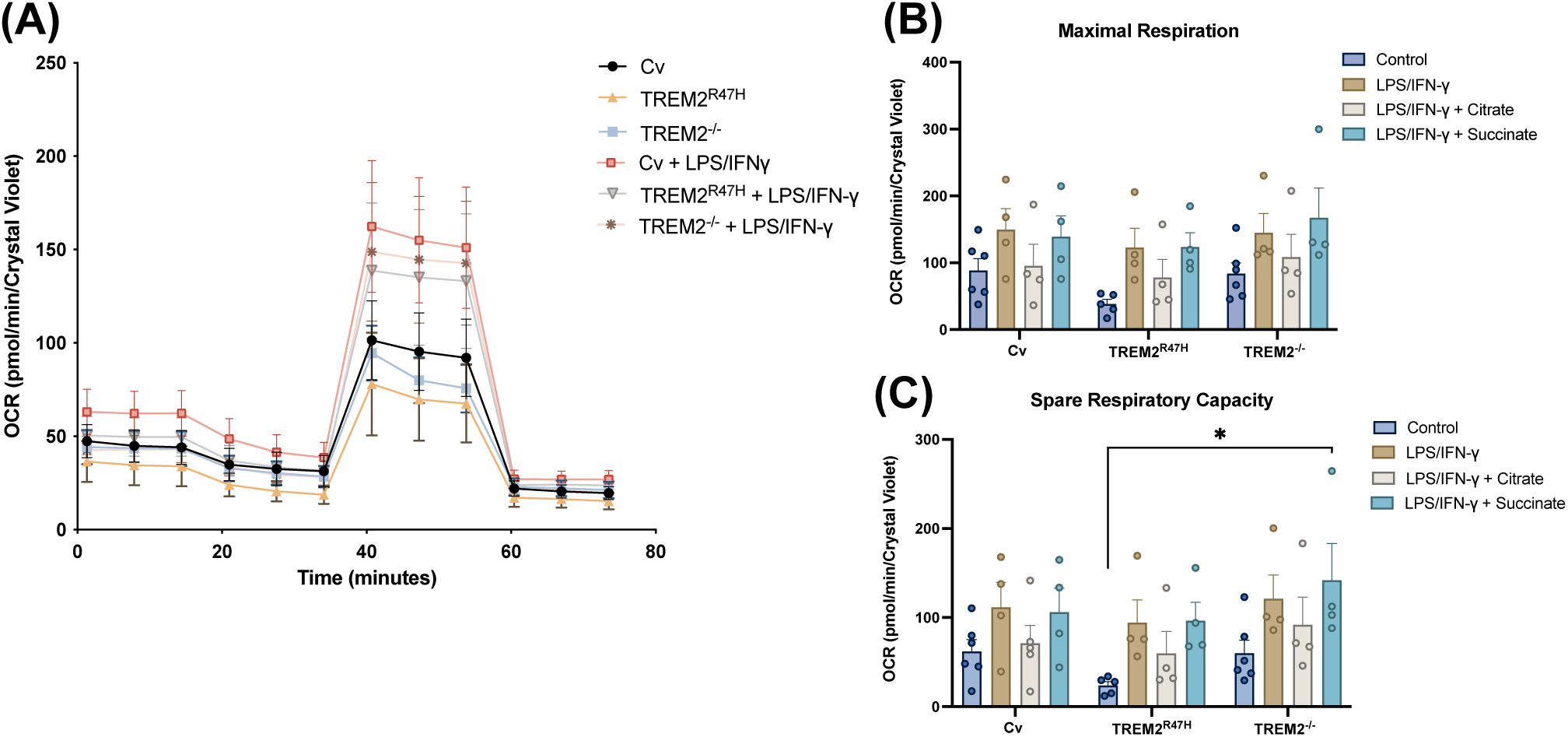
**(A)** Metabolic analysis of oxygen consumption rate (OCR) **(B)** Maximal respiration, and **(C)** spare respiratory capacity in Common variant (Cv), TREM2^R47H^ and TREM2^−/−^ iPS-Mg after 24 h LPS/IFN-γ stimulation and treatment with succinate (10 mM) or citrate (10 mM). Data are presented as mean ± SEM. Statistical significance was addressed using One-way ANOVA or Two-way ANOVA with Tukey post-hoc analysis to compare controls and treated groups respectively; *p<0.05 (n=5-6)

**Supplementary Figure 2.**
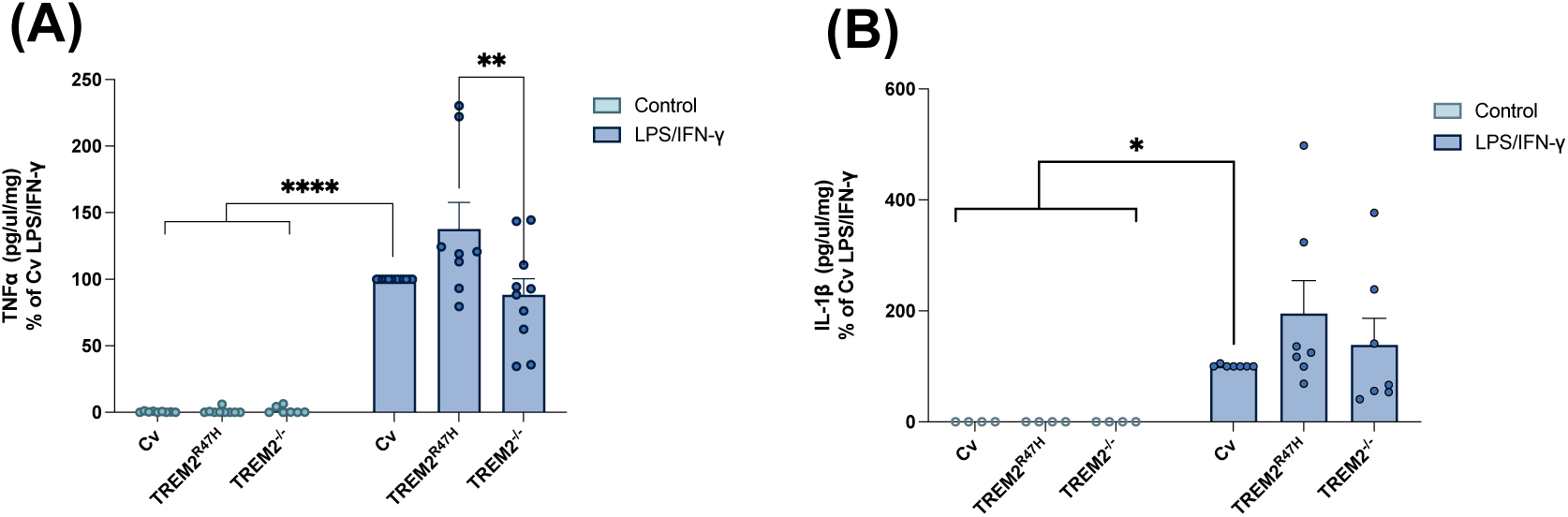
**(A)** Basal and LPS/IFN-γ evoked pro-inflammatory release of TNFα and **(B)** IL-1β were assessed by ELISA in the culture medium of iPS-Mg expressing common variant TREM2, TREM2^R47H^ and TREM2^−/−^. Data are presented as mean ± SEM. Statistical significance was addressed using two-way ANOVA with Tukey post-hoc analysis *p<0.05 (n=4-10)

## Notes

### Competing Interest Statement

The authors have declared no competing interest.

## References

1. Guerreiro R, Wojtas A, Bras J, Carrasquillo M, Rogaeva E, Majounie E, Cruchaga C, Sassi C, Kauwe JSK, Younkin S, et al. TREM2 variants in Alzheimer’s disease. N Engl J Med (2013) 368:117–127. doi: 10.1056/NEJMoa1211851

2. Jonsson T, Stefansson H, Steinberg S, Jonsdottir I, Jonsson PV, Snaedal J, Bjornsson S, Huttenlocher J, Levey AI, Lah JJ, et al. Variant of TREM2 associated with the risk of Alzheimer’s disease. N Engl J Med (2013) 368:107–116. doi: 10.1056/NEJMoa1211103

3. Wang Y, Ulland TK, Ulrich JD, Song W, Tzaferis JA, Hole JT, Yuan P, Mahan TE, Shi Y, Gilfillan S, et al. TREM2-mediated early microglial response limits diffusion and toxicity of amyloid plaques. J Exp Med (2016) 213:667–675. doi: 10.1084/jem.20151948

4. Ulrich JD, Finn MB, Wang Y, Shen A, Mahan TE, Jiang H, Stewart FR, Piccio L, Colonna M, Holtzman DM. Altered microglial response to Aβ plaques in APPPS1-21 mice heterozygous for TREM2. Mol Neurodegeneration (2014) 9:20. doi: 10.1186/1750-1326-9-20

5. Yeh FL, Wang Y, Tom I, Gonzalez LC, Sheng M. TREM2 Binds to Apolipoproteins, Including APOE and CLU/APOJ, and Thereby Facilitates Uptake of Amyloid-Beta by Microglia. Neuron (2016) 91:328–340. doi: 10.1016/j.neuron.2016.06.015

6. Ulland TK, Song WM, Huang SC-C, Ulrich JD, Sergushichev A, Beatty WL, Loboda AA, Zhou Y, Cairns NJ, Kambal A, et al. TREM2 Maintains Microglial Metabolic Fitness in Alzheimer’s Disease. Cell (2017) 170:649–663.e13. doi: 10.1016/j.cell.2017.07.023

7. Orihuela R, McPherson CA, Harry GJ. Microglial M1/M2 polarization and metabolic states. British J Pharmacology (2016) 173:649–665. doi: 10.1111/bph.13139

8. Lynch MA. Can the emerging field of immunometabolism provide insights into neuroinflammation? Progress in Neurobiology (2020) 184:101719. doi: 10.1016/j.pneurobio.2019.101719

9. Van den Bossche J, Baardman J, Otto NA, van der Velden S, Neele AE, van den Berg SM, Luque-Martin R, Chen H-J, Boshuizen MCS, Ahmed M, et al. Mitochondrial Dysfunction Prevents Repolarization of Inflammatory Macrophages. Cell Reports (2016) 17:684–696. doi: 10.1016/j.celrep.2016.09.008

10. Garcia-Reitboeck P, Phillips A, Piers TM, Villegas-Llerena C, Butler M, Mallach A, Rodrigues C, Arber CE, Heslegrave A, Zetterberg H, et al. Human Induced Pluripotent Stem Cell-Derived Microglia-Like Cells Harboring TREM2 Missense Mutations Show Specific Deficits in Phagocytosis. Cell Reports (2018) 24:2300–2311. doi: 10.1016/j.celrep.2018.07.094

11. Piers TM, Cosker K, Mallach A, Johnson GT, Guerreiro R, Hardy J, Pocock JM. A locked immunometabolic switch underlies TREM2 R47H loss of function in human iPSC-derived microglia. FASEB j (2020) 34:2436–2450. doi: 10.1096/fj.201902447R

12. Cosker K, Mallach A, Limaye J, Piers TM, Staddon J, Neame SJ, Hardy J, Pocock JM. Microglial signalling pathway deficits associated with the patient derived R47H TREM2 variants linked to AD indicate inability to activate inflammasome. Sci Rep (2021) 11:13316. doi: 10.1038/s41598-021-91207-1

13. Mallach A, Gobom J, Arber C, Piers TM, Hardy J, Wray S, Zetterberg H, Pocock J. Differential Stimulation of Pluripotent Stem Cell-Derived Human Microglia Leads to Exosomal Proteomic Changes Affecting Neurons. Cells (2021) 10:2866. doi: 10.3390/cells10112866

14. Mallach A, Gobom J, Zetterberg H, Hardy J, Piers TM, Wray S, Pocock JM. The influence of the R47H triggering receptor expressed on myeloid cells 2 variant on microglial exosome profiles. Brain Communications (2021) 3:fcab009. doi: 10.1093/braincomms/fcab009

15. Wang L, Zhang Y, Kiprowska M, Guo Y, Yamamoto K, Li X. Diethyl Succinate Modulates Microglial Polarization and Activation by Reducing Mitochondrial Fission and Cellular ROS. Metabolites (2021) 11:854. doi: 10.3390/metabo11120854

16. Wang Q, Lu M, Zhu X, Gu X, Zhang T, Xia C, Yang L, Xu Y, Zhou M. The role of microglia immunometabolism in neurodegeneration: Focus on molecular determinants and metabolic intermediates of metabolic reprogramming. Biomedicine & Pharmacotherapy (2022) 153:113412. doi: 10.1016/j.biopha.2022.113412

17. Bernier L-P, York EM, Kamyabi A, Choi HB, Weilinger NL, MacVicar BA. Microglial metabolic flexibility supports immune surveillance of the brain parenchyma. Nat Commun (2020) 11:1559. doi: 10.1038/s41467-020-15267-z

18. Fairley LH, Wong JH, Barron AM. Mitochondrial Regulation of Microglial Immunometabolism in Alzheimer’s Disease. Front Immunol (2021) 12:624538. doi: 10.3389/fimmu.2021.624538

19. Nair S, Sobotka KS, Joshi P, Gressens P, Fleiss B, Thornton C, Mallard C, Hagberg H. Lipopolysaccharide-induced alteration of mitochondrial morphology induces a metabolic shift in microglia modulating the inflammatory response in vitro and in vivo. Glia (2019) 67:1047–1061. doi: 10.1002/glia.23587

20. Jha NK, Jha SK, Kar R, Nand P, Swati K, Goswami VK. Nuclear factor-kappa β as a therapeutic target for Alzheimer’s disease. Journal of Neurochemistry (2019) 150:113–137. doi: 10.1111/jnc.14687

21. Xiang X, Piers TM, Wefers B, Zhu K, Mallach A, Brunner B, Kleinberger G, Song W, Colonna M, Herms J, et al. The Trem2 R47H Alzheimer’s risk variant impairs splicing and reduces Trem2 mRNA and protein in mice but not in humans. Mol Neurodegeneration (2018) 13:49. doi: 10.1186/s13024-018-0280-6

22. Damisah EC, Rai A, Grutzendler J. TREM2: Modulator of Lipid Metabolism in Microglia. Neuron (2020) 105:759–761. doi: 10.1016/j.neuron.2020.02.008

23. Monsorno K, Buckinx A, Paolicelli RC. Microglial metabolic flexibility: emerging roles for lactate. Trends in Endocrinology & Metabolism (2022) 33:186–195. doi: 10.1016/j.tem.2021.12.001

24. Zhao Y, Xu H. Microglial lactate metabolism as a potential therapeutic target for Alzheimer’s disease. Mol Neurodegeneration (2022) 17:36. doi: 10.1186/s13024-022-00541-z

25. Garcia-Reitboeck P, Phillips A, Piers TM, Villegas-Llerena C, Butler M, Mallach A, Rodrigues C, Arber CE, Heslegrave A, Zetterberg H, et al. Human Induced Pluripotent Stem Cell-Derived Microglia-Like Cells Harboring TREM2 Missense Mutations Show Specific Deficits in Phagocytosis. Cell Reports (2018) 24:2300–2311. doi: 10.1016/j.celrep.2018.07.094

26. Wang W-Y, Tan M-S, Yu J-T, Tan L. Role of pro-inflammatory cytokines released from microglia in Alzheimer’s disease. Ann Transl Med (2015) 3:136. doi: 10.3978/j.issn.2305-5839.2015.03.49

27. Su F, Bai F, Zhang Z. Inflammatory Cytokines and Alzheimer’s Disease: A Review from the Perspective of Genetic Polymorphisms. Neurosci Bull (2016) 32:469–480. doi: 10.1007/s12264-016-0055-4

28. Tannahill GM, Curtis AM, Adamik J, Palsson-McDermott EM, McGettrick AF, Goel G, Frezza C, Bernard NJ, Kelly B, Foley NH, et al. Succinate is an inflammatory signal that induces IL-1β through HIF-1α. Nature (2013) 496:238–242. doi: 10.1038/nature11986

29. Infantino V, Convertini P, Cucci L, Panaro MA, Di Noia MA, Calvello R, Palmieri F, Iacobazzi V. The mitochondrial citrate carrier: a new player in inflammation. Biochemical Journal (2011) 438:433–436. doi: 10.1042/BJ20111275

30. Williams NC, O’Neill LAJ. A Role for the Krebs Cycle Intermediate Citrate in Metabolic Reprogramming in Innate Immunity and Inflammation. Front Immunol (2018) 9:141. doi: 10.3389/fimmu.2018.00141

31. Lu R-C, Tan M-S, Wang H, Xie A-M, Yu J-T, Tan L. Heat Shock Protein 70 in Alzheimer’s Disease. BioMed Research International (2014) 2014:1–8. doi: 10.1155/2014/435203

32. Sun E, Motolani A, Campos L, Lu T. The Pivotal Role of NF-kB in the Pathogenesis and Therapeutics of Alzheimer’s Disease. IJMS (2022) 23:8972. doi: 10.3390/ijms23168972

33. Wang Y, Cella M, Mallinson K, Ulrich JD, Young KL, Robinette ML, Gilfillan S, Krishnan GM, Sudhakar S, Zinselmeyer BH, et al. TREM2 Lipid Sensing Sustains the Microglial Response in an Alzheimer’s Disease Model. Cell (2015) 160:1061–1071. doi: 10.1016/j.cell.2015.01.049

34. Nugent AA, Lin K, Van Lengerich B, Lianoglou S, Przybyla L, Davis SS, Llapashtica C, Wang J, Kim DJ, Xia D, et al. TREM2 Regulates Microglial Cholesterol Metabolism upon Chronic Phagocytic Challenge. Neuron (2020) 105:837–854.e9. doi: 10.1016/j.neuron.2019.12.007

35. Ulland TK, Song WM, Huang SC-C, Ulrich JD, Sergushichev A, Beatty WL, Loboda AA, Zhou Y, Cairns NJ, Kambal A, et al. TREM2 Maintains Microglial Metabolic Fitness in Alzheimer’s Disease. Cell (2017) 170:649–663.e13. doi: 10.1016/j.cell.2017.07.023

36. Jha AK, Huang SC-C, Sergushichev A, Lampropoulou V, Ivanova Y, Loginicheva E, Chmielewski K, Stewart KM, Ashall J, Everts B, et al. Network Integration of Parallel Metabolic and Transcriptional Data Reveals Metabolic Modules that Regulate Macrophage Polarization. Immunity (2015) 42:419–430. doi: 10.1016/j.immuni.2015.02.005

37. Kaneko I, Yamada N, Sakuraba Y, Kamenosono M, Tutumi S. Suppression of Mitochondrial Succinate Dehydrogenase, a Primary Target of β-Amyloid, and Its Derivative Racemized at Ser Residue. Journal of Neurochemistry (1995) 65:2585–2593. doi: 10.1046/j.1471-4159.1995.65062585.x

38. Williams NC, O’Neill LAJ. A Role for the Krebs Cycle Intermediate Citrate in Metabolic Reprogramming in Innate Immunity and Inflammation. Front Immunol (2018) 9:141. doi: 10.3389/fimmu.2018.00141

39. Mills E, O’Neill LAJ. Succinate: a metabolic signal in inflammation. Trends in Cell Biology (2014) 24:313–320. doi: 10.1016/j.tcb.2013.11.008

40. Giorgi-Coll S, Amaral AI, Hutchinson PJA, Kotter MR, Carpenter KLH. Succinate supplementation improves metabolic performance of mixed glial cell cultures with mitochondrial dysfunction. Sci Rep (2017) 7:1003. doi: 10.1038/s41598-017-01149-w

41. Jalloh I, Helmy A, Howe DJ, Shannon RJ, Grice P, Mason A, Gallagher CN, Stovell MG, Van Der Heide S, Murphy MP, et al. Focally perfused succinate potentiates brain metabolism in head injury patients. J Cereb Blood Flow Metab (2017) 37:2626–2638. doi: 10.1177/0271678X16672665

42. Dikalov S. Cross talk between mitochondria and NADPH oxidases. Free Radical Biology and Medicine (2011) 51:1289–1301. doi: 10.1016/j.freeradbiomed.2011.06.033

43. Mills EL, Kelly B, Logan A, Costa ASH, Varma M, Bryant CE, Tourlomousis P, Däbritz JHM, Gottlieb E, Latorre I, et al. Succinate Dehydrogenase Supports Metabolic Repurposing of Mitochondria to Drive Inflammatory Macrophages. Cell (2016) 167:457–470.e13. doi: 10.1016/j.cell.2016.08.064

44. Atagi Y, Liu C-C, Painter MM, Chen X-F, Verbeeck C, Zheng H, Li X, Rademakers R, Kang SS, Xu H, et al. Apolipoprotein E Is a Ligand for Triggering Receptor Expressed on Myeloid Cells 2 (TREM2). Journal of Biological Chemistry (2015) 290:26043–26050. doi: 10.1074/jbc.M115.679043

45. Bailey CC, DeVaux LB, Farzan M. The Triggering Receptor Expressed on Myeloid Cells 2 Binds Apolipoprotein E. Journal of Biological Chemistry (2015) 290:26033–26042. doi: 10.1074/jbc.M115.677286

46. Sudom A, Talreja S, Danao J, Bragg E, Kegel R, Min X, Richardson J, Zhang Z, Sharkov N, Marcora E, et al. Molecular basis for the loss-of-function effects of the Alzheimer’s disease–associated R47H variant of the immune receptor TREM2. Journal of Biological Chemistry (2018) 293:12634–12646. doi: 10.1074/jbc.RA118.002352

47. Ulland TK, Colonna M. TREM2 — a key player in microglial biology and Alzheimer disease. Nat Rev Neurol (2018) 14:667–675. doi: 10.1038/s41582-018-0072-1

48. Wang Y, Cella M, Mallinson K, Ulrich JD, Young KL, Robinette ML, Gilfillan S, Krishnan GM, Sudhakar S, Zinselmeyer BH, et al. TREM2 Lipid Sensing Sustains the Microglial Response in an Alzheimer’s Disease Model. Cell (2015) 160:1061–1071. doi: 10.1016/j.cell.2015.01.049

49. Wei W, Zhang L, Xin W, Pan Y, Tatenhorst L, Hao Z, Gerner ST, Huber S, Juenemann M, Butz M, et al. TREM2 regulates microglial lipid droplet formation and represses post-ischemic brain injury. Biomedicine & Pharmacotherapy (2024) 170:115962. doi: 10.1016/j.biopha.2023.115962

50. Filipello F, You S-F, Mirfakhar FS, Mahali S, Bollman B, Acquarone M, Korvatska O, Marsh JA, Sivaraman A, Martinez R, et al. Defects in lysosomal function and lipid metabolism in human microglia harboring a TREM2 loss of function mutation. Acta Neuropathol (2023) 145:749–772. doi: 10.1007/s00401-023-02568-y

51. Gimeno-Bayón J, López-López A, Rodríguez MJ, Mahy N. Glucose pathways adaptation supports acquisition of activated microglia phenotype. J of Neuroscience Research (2014) 92:723–731. doi: 10.1002/jnr.23356

52. Voloboueva LA, Emery JF, Sun X, Giffard RG. Inflammatory response of microglial BV-2 cells includes a glycolytic shift and is modulated by mitochondrial glucose-regulated protein 75/mortalin. FEBS Letters (2013) 587:756–762. doi: 10.1016/j.febslet.2013.01.067

53. Pan R-Y, He L, Zhang J, Liu X, Liao Y, Gao J, Liao Y, Yan Y, Li Q, Zhou X, et al. Positive feedback regulation of microglial glucose metabolism by histone H4 lysine 12 lactylation in Alzheimer’s disease. Cell Metabolism (2022) 34:634–648.e6. doi: 10.1016/j.cmet.2022.02.013

54. Cheng J, Zhang R, Xu Z, Ke Y, Sun R, Yang H, Zhang X, Zhen X, Zheng L-T. Early glycolytic reprogramming controls microglial inflammatory activation. J Neuroinflammation (2021) 18:129. doi: 10.1186/s12974-021-02187-y

55. Fodelianaki G, Lansing F, Bhattarai P, Troullinaki M, Zeballos MA, Charalampopoulos I, Gravanis A, Mirtschink P, Chavakis T, Alexaki VI. Nerve Growth Factor modulates LPS - induced microglial glycolysis and inflammatory responses. Experimental Cell Research (2019) 377:10–16. doi: 10.1016/j.yexcr.2019.02.023

56. Shi J, Fan J, Su Q, Yang Z. Cytokines and Abnormal Glucose and Lipid Metabolism. Front Endocrinol (2019) 10:703. doi: 10.3389/fendo.2019.00703

57. Hu R, Xia C-Q, Butfiloski E, Clare-Salzler M. Effect of high glucose on cytokine production by human peripheral blood immune cells and type I interferon signaling in monocytes: Implications for the role of hyperglycemia in the diabetes inflammatory process and host defense against infection. Clinical Immunology (2018) 195:139–148. doi: 10.1016/j.clim.2018.06.003

58. Zhong L, Xu Y, Zhuo R, Wang T, Wang K, Huang R, Wang D, Gao Y, Zhu Y, Sheng X, et al. Soluble TREM2 ameliorates pathological phenotypes by modulating microglial functions in an Alzheimer’s disease model. Nat Commun (2019) 10:1365. doi: 10.1038/s41467-019-09118-9

59. Liu W, Taso O, Wang R, Bayram S, Graham AC, Garcia-Reitboeck P, Mallach A, Andrews WD, Piers TM, Botia JA, et al. *Trem2* promotes anti-inflammatory responses in microglia and is suppressed under pro-inflammatory conditions. Human Molecular Genetics (2020) 29:3224–3248. doi: 10.1093/hmg/ddaa209

60. Biel D, Suárez-Calvet M, Hager P, Rubinski A, Dewenter A, Steward A, Roemer S, Ewers M, Haass C, Brendel M, et al. STREM2 is associated with amyloid-related p-tau increases and glucose hypermetabolism in Alzheimer’s disease. EMBO Mol Med (2023) 15:e16987. doi: 10.15252/emmm.202216987

61. Zhang X, Tang L, Yang J, Meng L, Chen J, Zhou L, Wang J, Xiong M, Zhang Z. Soluble TREM2 ameliorates tau phosphorylation and cognitive deficits through activating transgelin-2 in Alzheimer’s disease. Nat Commun (2023) 14:6670. doi: 10.1038/s41467-023-42505-x

