## Supplementary figures and images for "Amelioration of signalling deficits underlying metabolic shortfall in TREM2^R47H^ human iPSC-derived microglia"

### Supplementary Figures 1 and 2

#### Supplementary figure 1

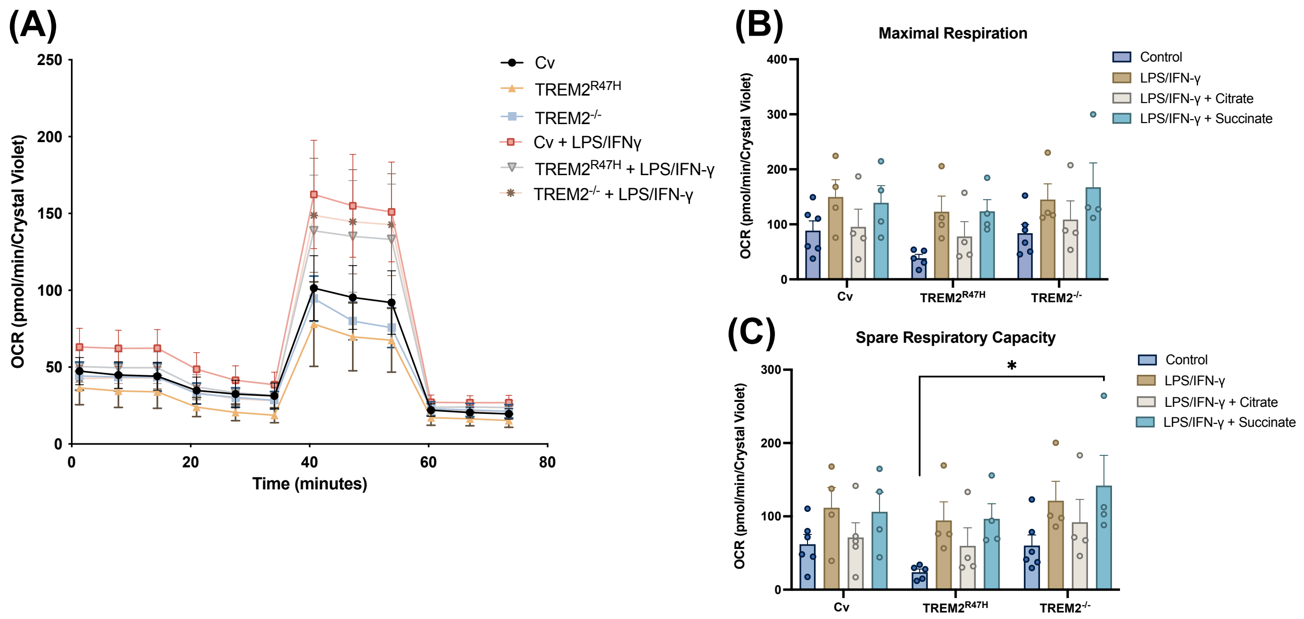


#### Supplementary figure 2

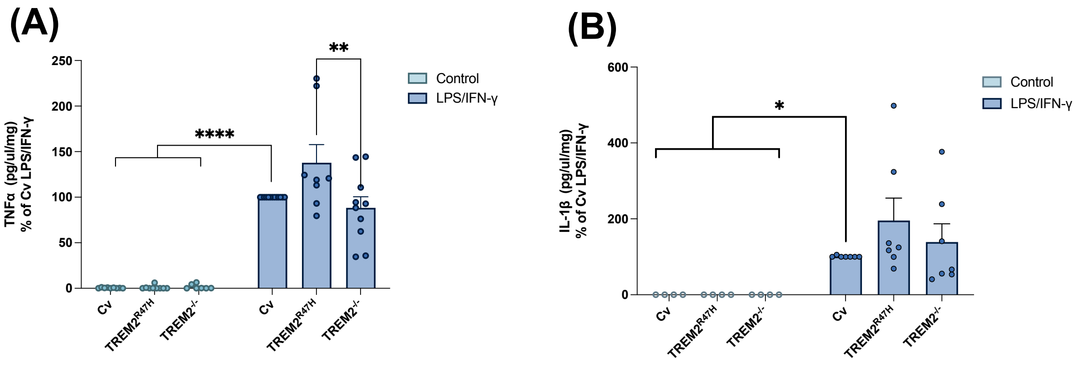
